# Bioremediation Potential of Select Bacterial Species for the Neonicotinoid Insecticides, Thiamethoxam and Imidacloprid

**DOI:** 10.1101/2020.07.26.221945

**Authors:** Stephanie M. Zamule, Cassandra Dupre, Meghan Mendola, Julia Widmer, Jane Shebert, Carol E. Roote, Padmini Das

## Abstract

The neonicotinoid insecticides, including thiamethoxam (THM) and imidacloprid (IMI), have become increasingly favored in the past decade due to their specificity as insect neurotoxicants. However, neonicotinoids have been implicated as a potential contributing factor in Colony Collapse Disorder (CCD), the widespread disappearance of honey bees, which affects produce production on a global scale. The environmental persistence of neonicotinoids underscores the importance of developing a sustainable, ecologically-friendly remediation technique to remove residual insecticides from the environment. The present study characterizes the neonicotinoid bioremediation potential of six bacterial species: *Pseudomonas fluorescens, Pseudomonas putida, Pseudomonas aeruginosa, Alcaligenes faecalis, Escherichia coli*, and *Streptococcus lactis*. In Phase I, we evaluated the utilization of IMI or THM as the sole carbon or nitrogen source by *P. fluorescens, P. putida*, and *P. aeruginosa*. All three species were better able to utilize THM over IMI as their sole carbon or nitrogen source, and better growth was noted when THM was used as the sole nitrogen source compared to the sole carbon source. Thus, further studies proceeded with THM only. In Phase II, we assessed the kinetics of THM removal from aqueous media by the six species. Significant (p<0.0001) reductions in 70 mg/L THM concentration were observed for *P. fluorescens* (67%), *P. putida* (65%), *P. aeruginosa* (52%), and *A. faecalis* (39%) over the 24-day study period, and for *E. coli* (60%) and *S. lactis* (12%) over the 14-day study period. The amount of time required to remove 50% of the THM in the media (T_50_) was: 12 days (d) (*E. coli*), 18 d (*P. fluorescens*), 19 d (*P. putida*), and 23 d (*P. aeruginosa*). Neither *A. faecalis* nor *S. lactis* achieved 50% removal during the study periods. The THM removal by all species followed a first-order kinetic reaction and half-lives were calculated accordingly. HPLC chromatograms of *P. fluorescens, P. putida*, and *E. coli* cultures revealed that as the area of the THM peak decreased over time, the area of an unidentified metabolite peak increased. In Phase II, we sought to characterize this metabolite and the overall metabolic efficiency of these three species. Maximal THM removal occurred at 30°C for all bacterial species assessed. Identification of the metabolite is currently underway, which will allow determination of whether the metabolite is less toxic than the parent compound, a prerequisite for this remediation technique to be viable. If the metabolite is found to be less hazardous than THM, further testing will follow to evaluate the use of this bioremediation technique in the field.

## Introduction

Neonicotinoid pesticides, including thiamethoxam (THM) and imidacloprid (IMI), have become increasingly popular in the last two decades due to their high specificity as insect neurotoxicants (Simon-Delso et al. 2015). As of 2009, IMI was registered for use on 140 crops with annual sales of US$1.09 billion worldwide (Simon-Delso et al. 2015). As of 2012, THM, the primary focus of this study, was registered for use on 115 different crops and generated US$1.1 billion in annual sales worldwide (Simon-Delso et al. 2015). While the specificity of neonicotinoids substantially reduces their toxicity to mammals, potential sublethal effects on off-target insects, including honey bees, is a concern (Fairbrother et al. 2014). Specifically, neonicotinoids are a potential contributing factor to Colony Collapse Disorder (CCD), a phenomena characterized by the disappearance of worker bees (Fairbrother et al. 2014). Because animal pollinators, including the honey bee, are required for production of approximately one-third of crop production globally (Klein et al. 2007), and animal-pollinated crops are worth an estimated $175 billion dollars per year globally (Gallai et al. 2009), CCD has substantial health, economic, and ecological implications worldwide. While a number of laboratory studies have demonstrated neonicotinoid toxicity in bees, whether these effects exist in the field at environmentally-relevant concentrations remains equivocal (Fairbrother et al. 2014, Carreck 2017). However, concern has been such that, as of May 2018, the European Union has banned the use of IMI, THM, and a third neonicotinoid, clothianidin, for all outdoor uses (European Commission 2019).

CCD is the disappearance of almost all adult bees from a hive, with no dead bees present (Kaplan 2012). There is not a single cause of CCD, but instead several broad stress factors that are believed to cause it. These include pathogens, parasites, environmental stressors such as pesticides, and hive management stressors (Kaplan 2012). IMI and THM both affect the survival of bees and their colonies. The average neonicotinoid concentrations in the field are 1.9 ng/g in nectar and 6.1 ng/g in pollen (Wood et al. 2018). The median lethal concentration of imidacloprid to bees is 1.4651 ng imidacloprid/uL nectar or sucrose solution with doses under that leading to changes in foraging patterns, flight endurance and immune system complications (Catae et al. 2017). IMI has been shown to change the expression of nicotinic acetylcholine receptor alpha 1 which is used in synapses of the brain, leading to possible inhibition of cognitive processes and memory building (Catae et al. 2017). Not only is cognition affected, but ability to effectively forage is decreased. IMI does not affect the bees physically per se, but instead increases flight velocity leading to a flight duration and distance that is a third of what an untreated bee is capable of (Kenna et al. 2019). This decrease in duration leads to a much smaller foraging area and a decreased abundance and diversity of food for the hive (Kenna et al. 2019). THM has a similar effect, causing excitement and increased velocity with negative effects of duration decreasing by 54% and distance decreasing by 56% when the honey bees are continuously exposed (Tosi et al. 2017). Exposure to THM affected bees’ ability to differentiate between floral scents and blank scents when in a lab run maze and caused the bees to become indecisive in deciding directions, causing longer durations to complete the maze (Jiang et al. 2018). Bees treated with IMI also show decreased motivation to forage and visit fewer flowers from the beginning and don’t visit them as often (Lamsa et al. 2018). 23% fewer bees return to collect nectar with IMI, and those that do return to collect nectar collect 63% less when IMI levels are at 40 ug/L (Tan et al. 2014). When treated with THM, the bees were less responsive to sugar which caused them to consume less than normal amounts (Jiang et al. 2018). When queen bees are exposed to IMI, nest initiation is delayed and brood numbers decrease, leading to a smaller colony and lessened activity which strains survival (Leza et al. 2018). THM exposure causes a decrease in colony size, reducing the hive cumulative weight gain of honey by 30.2% and reducing clusters by 21.7% (Wood et al. 2018). THM and IMI cause detrimental changes to bees and their colonies, leading to altered foraging and immunity, and reducing colony size and health.

Neonicotinoids persist in soil well beyond their agricultural usefulness, thus extending the time during which honey bees and other off-target species may come into contact with these compounds. For example, IMI has a half-life up to ~400 days depending on different environmental conditions such as soil composition and vegetation (Anhalt et al. 2008). THM has been shown to exhibit a half-life of up to ~300 days, depending on soil conditions (Gupta et al. 2008). These large half-lives, in addition to only 5% of the active neonicotinoid ingredients being taken up by the crops while the rest disperses into the surrounding environment, lead to an increasingly urgent environmental issue (Sur and Stork 2003). Bioremediation offers a potentially sustainable, inexpensive, ecologically-friendly solution to the problem of neonicotinoid persistence in the environment.

Bacterial species that have been shown to degrade IMI include a *Leifsonia* strain PC-21 (37-58% reduction of 25 mg/L over 21 days) (Anhalt et al. 2007), *Trichoderma viride* (50% reduction of 4 ppm over 12 days) (Tamilselvan et al. 2008), *Burkholderia cepacia* CH9 (69% reduction of 50 ppm over 20 days) (Madhuban et al. 2011), *Ochrobactrum anthropic* W-7 (67.67% reduction of 50 mg/L over 48 hours) (Hu et al. 2013), *Klebsiella pneumonia* BHC1 (78% reduction of 50 mg/L over 7 days) (Phugare et al. 2013), *Achromobacter* strain R-46660 (62% reduction of 100 ppm over 20 days), *Microbacterium* B-2013 (60% reduction of 100 ppm over 20 days) (Negi et al. 2014), *Rhizobium* sp. (45.48% reduction of 25 mg/L over 25 days), *Bacillus subtilis* (37.20% reduction of 25 mg/L over 25 days), *Brevibacterium* sp. (29.88% reduction of 25 mg/L over 25 days) (Sabourmoghaddam et al. 2014). Other species with IMI degrading ability, as measured by changes in chemical oxygen demand (COD) and biochemical oxygen demand (BOD_5_), include *Methylobacterium radiotolerans* and *Microbacterium arthosphaerae* (Erguven and Yildirim 2019) and *Ochrobactrum thiophenivorans* and *Sphingomonas melonis* (Erguven and Demirci 2019).

A number of groups have investigated the ability of various *Pseudomonas* species to degrade IMI. Tamilselvan et al. (2008) isolated *Pseudomonas fluorescens* and found this species was able to degrade 100% of 2 ppm IMI over 12 days, but only about 60% of 4 ppm IMI over that same time period. Pandey et al. (2009) identified three strains of *Pseudomonas* (1G, 1W, and GP2), each of which degraded approximately 70% of 50 ppm IMI within 14 days. Negi et al. (2014) isolated bacteria identified as *Pseudomonas* strain HY8N (GB35) that was capable of degrading 64% of 100 ppm IMI over 20 days. Sabourmoghaddam et al. (2014) identified a strain of *Pseudomonas putida* F1 able to degrade 32.4% of 25 mg/L IMI over 25 days. Most recently, Gupta et al. (2016) isolated *Pseudomonas* RPT 52 and found that this species degraded 46.5% of 0.5 mM IMI over 40 hours.

Research evaluating bacterial species capable of degrading THM is currently more limited (for review, see Hussain et al. 2016). Species that have been shown to degrade THM include: *Bacillus subtilis* FZB24, *Bacillus amyloliquefaciens* IN937a, *Bacillus pumilus* SE34 (ranging from 11-22% reduction of 0.25-2.5 mg/L IMI over 3 days) (Myresiotis et al. 2012); *Ensifer adhaerens* TXM-23 (37% reduction of 200 mg/L over 25 days) (Zhou et al. 2013), and a number of other members of the *Bacillus* genus, most notably *Bacillus aerophilus* IMBL 4.1 (45% reduction of 50 ug/mL over 15 days) (Rana et al. 2015). Additionally, Zhou et al. (2014) identified a soil enrichment culture comprised of members of the genera *Achromobacter, Agromyces, Ensifer, Mesorhizobium, Microbacterium* and *Pseudoxanthomonas t*hat was able to degrade 96% of 200 mg/L THM over 30 days.

A number of *Pseudomonas* species have emerged as perhaps the most promising candidates thus far for THM degradation. Pandey et al. (2009) identified three members of the *Pseudomonas* genus (strains 1G, 1W, and GP2), each capable of reducing 50 ppm THM in aqueous media by ~70% over 14 days. Rana et al. (2015) identified three *Pseudomonas* strains capable of reducing 50 ug/mL THM in aqueous media over 15 days: *Pseudomonas fulva* (IMBL 5.1) led to a 32% reduction, *Pseudomonas monteilii* (IMBL 4.3) led to a 34% reduction, and *Pseudomonas putida* (IMBL 5.2) led to a 38% reduction. In both aforementioned studies, culture media contained additional carbon and nitrogen sources, and thus THM did not serve as the sole source of these elements (Pandey et al. 2009, Rana et al. 2015). However, Rana et al. (2015) further found that THM reduction by *Pseudomonas putida* (IMBL 5.2) could be increased to 51% if THM served as the sole carbon and nitrogen source for the organism, although overall growth was reduced. However, these conditions would be challenging to simulate in the field. Thus, there is a clear need for further research in this area, both to more fully characterize the THM-degrading ability of *Pseudomonas* species and to identify additional species with THM bioremediative potential.

The present study assesses the ability of three *Pseudomonas* species (*P. fluorescens, P. putida*, and *P. aeruginosa*), as well as *Alcaligenes faecalis, Escherichia coli*, and *Streptococcus lactis* to grow on media containing IMI or THM as the sole carbon or nitrogen source and characterizes the THM degradation capacities of these organisms in aqueous media under laboratory conditions. This work builds upon understanding of the substantial THM degradation abilities of *Pseudomonas* species, documents the increase in concentration of a metabolite that correlates with decreasing THM levels, and to our knowledge, is the first to report the significant THM degradation capacity of *E. coli*.

## Materials & Methods

### Chemicals and Organisms

Analytical grade thiamethoxam, imidacloprid, clothianidin, N-desmethylthiamethoxam, and nitroguanidine were purchased from Sigma-Aldrich (St. Louis, MO). *Pseudomonas putida* and *Pseudomonas fluorescens* were purchased from Ward’s Science (Rochester, NY). *Pseudomonas aeruginosa, Alcaligenes faecalis, Streptococcus lactis*, and *Escherichia coli* were purchased from Presque Isle Cultures (Erie, PA). 25mm syringe filters with 0.45 μm PTFE membranes were purchased from VWR International (Radnor, PA).

### Media and Culture Conditions

All chemical stocks were prepared by dissolving the compounds in sterilized deionized water. *Pseudomonas* minimal media (PMM) was prepared as follows: 98.5 mL distilled water, 1.5 mL glycerol, 0. 449 g ammonia sulfate, 0.15 g dibasic potassium phosphate, and 0.02 g magnesium sulfate. PMM with IMI or THM as the sole carbon source was prepared by substituting IMI or THM, respectively, for glycerol. PMM with IMI or THM as the sole nitrogen source was prepared by substituting IMI or THM, respectively, for ammonia sulfate. Bacteria were grown under non-shaking, aerobic conditions at the specified temperatures. Growth was assessed spectrophotometrically using a Thermo Scientific Genesys 20 by measuring optical density at 600 nm.

### High Performance Liquid Chromatography

Thiamethoxam depletion and metabolite levels were assessed using HPLC. Samples were prepared for analysis by centrifuging 1 ml aliquots at 9615 x g for 10 minutes. The supernatant was then removed and filtered using a 25 mm syringe filter with a 0.45 μm PTFE membrane. 5 μl of each sample was placed into a HPLC vial for analysis. The Agilent 1260 Infinity HPLC was used for analysis with a Supercoil LC-18DB 15 cm 4.6 mm (ID), 5 μm particle size column as a stationary phase. The mobile phase consisted of 70% deionized water and 30% acetonitrile with a flow rate of 1.0 mL/min. The diode array detector was set at a wavelength of 252 nm. Standard curves were made daily ranging from 0 mg/L to 160 mg/L of THM allowing for a comparison to unknown THM concentration. Unknown metabolites were compared to daily standard curves ranging from 0 mg/L to 160 mg/L of clothianidin, N-desmethylthiamethoxam and nitroguanine.

### Statistics and Kinetic Data Analysis

Residual THM data was fit to first order and second order kinetic models, and rate constants were calculated as a function of bacterial type and compared at 95% confidence intervals. All statistical tests were carried out using JMP Pro13. Two way ANOVA followed by mean comparison was carried out with the Tukey Kramer Honest Significant Difference (HSD) test.

## Results

*P. aeruginosa, P. putida*, and *P. fluorescens* were inoculated into PMM with IMI or THM as the sole source of carbon or nitrogen and grown under standard conditions. All three of these *Pseudomonas* species exhibited substantially more growth in media when THM served as the sole carbon or nitrogen source compared to when IMI served as the sole carbon or nitrogen source, although all experimental conditions showed less growth than controls grown in standard PMM (Fig. 1). In general, growth in media in which IMI or THM served as the sole nitrogen source was greater than growth in media in which the same compound served as the sole carbon source (Fig. 1). Compared to the other species assessed, *P. putida* grown in media in which THM served as the sole nitrogen source exhibited the most growth relative to controls (>62% of control growth for the 10 mg/L THM concentration) (Supplementary Data Table S1). Interestingly, for media in which THM served as the sole carbon or nitrogen source, growth appeared to be only nominally impacted by THM concentration (Fig. 1).

**FIGURE 1.**
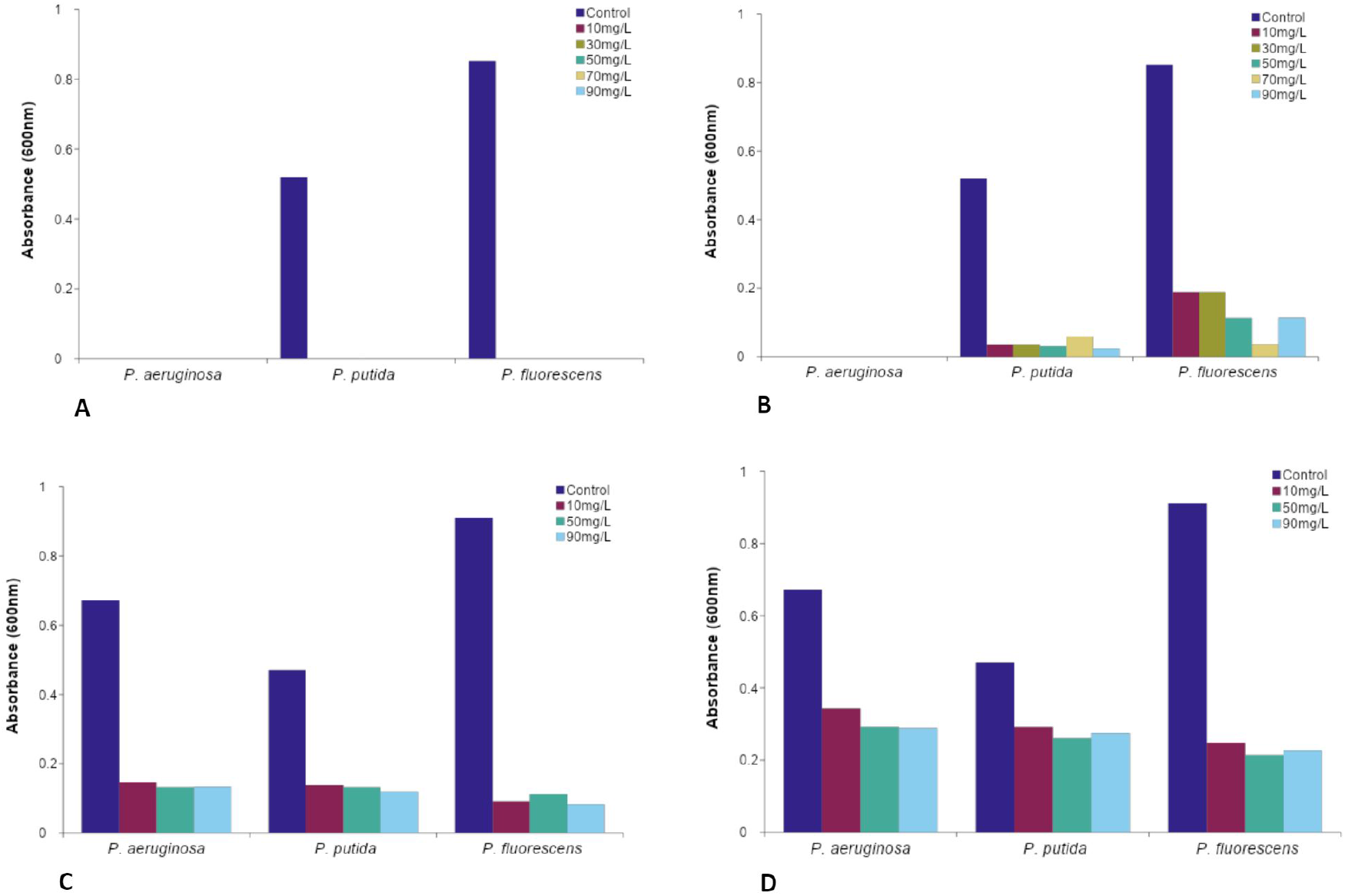
Optimization of *P. aeruginosa, P. putida*, and *P. fluorescens* growth conditions using imidacloprid and thiamethoxam as sole carbon or nitrogen source. Bacteria were cultured under standard aerobic conditions at 30°C in PMM with (A) imidacloprid as sole carbon source, (B) imidacloprid as sole nitrogen source, (C) thiamethoxam as sole carbon source, and (D) thiamethoxam as sole nitrogen source. Bacterial growth was measured by spectrophotometry at 600 nm after 5 days for imidacloprid and 21 days for thiamethoxam. Control = PMM inoculated with the species indicated.

Because growth of all three species assessed was greater in THM-substituted media than IMI-substituted media (Fig. 1), and additional studies showed a lack of growth of organisms in imidacloprid (data not shown), further studies focused on THM. In the next set of experiments, nutrient broth containing both a carbon and nitrogen source was used at only half-strength in an effort to push the organisms toward utilization of the THM in the media. When cultured in half-strength nutrient broth supplemented with 70 mg/L THM, significant (p<0.0001) reductions in THM concentration were observed for *P. fluorescens* (67%), *P. putida* (65%), *P. aeruginosa* (52%), and *A. faecalis* (39%) at the conclusion of the 24-day study period (Fig. 2). THM removal by these four species was significantly higher than that of the uninoculated control, confirming biotransformation of THM. All species exhibited a biphasic pattern of THM removal, initially slow and then steep, which showed best fit to the first order kinetics for these four species (see R^2^ for both first and second order fit in table 2). The first order reaction rate constants (-*k_1_*) for THM-removal increased with increasing time (Table 1). *P. fluorescens* and *P. putida* showed significantly higher removal rate constants after 24 days as compared to those of *P. aeruginosa* and *A. faecalis*.

**FIGURE 2.**
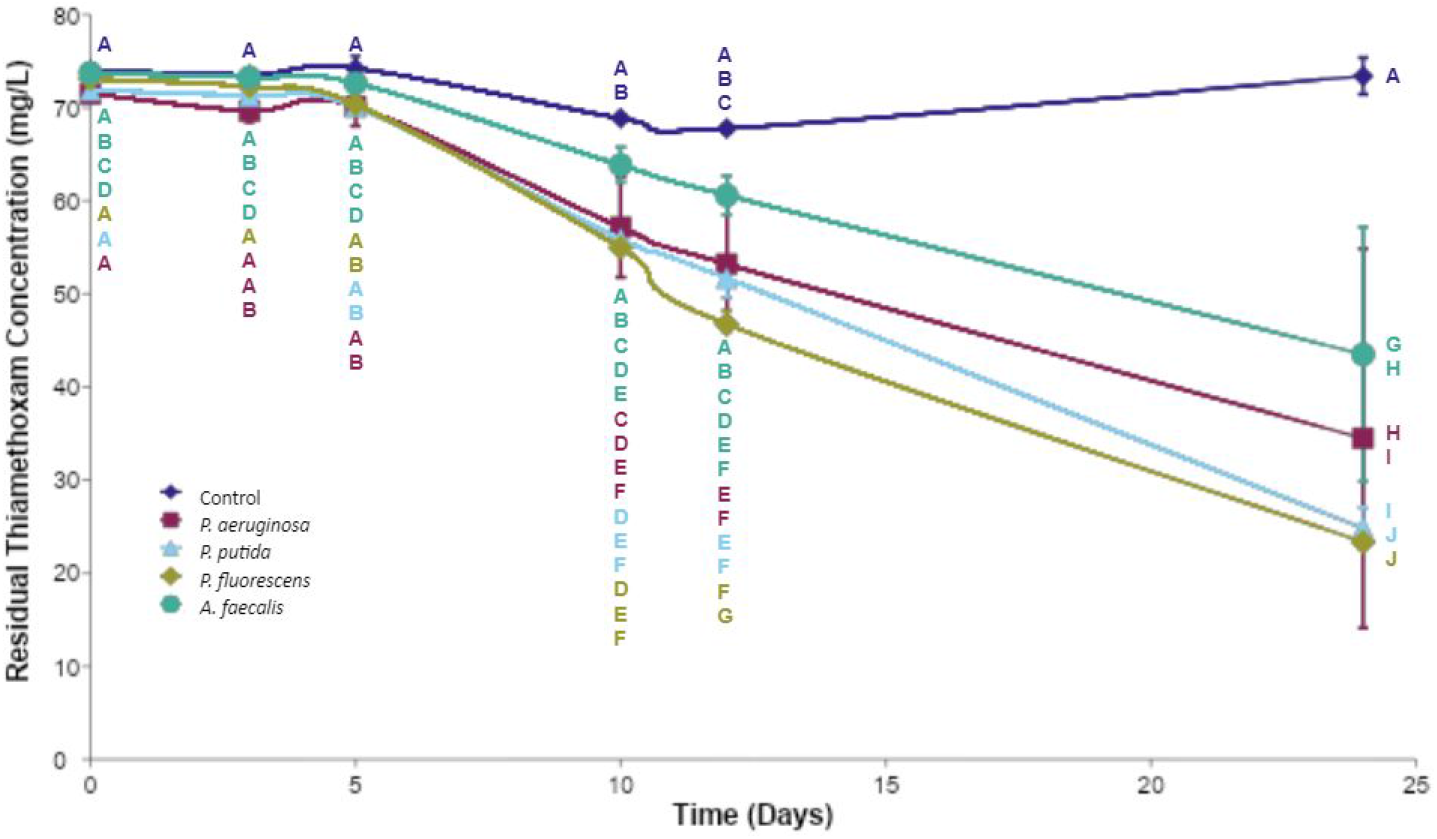
Removal of thiamethoxam from aqueous solution by *P. aeruginosa, P. putida, P. fluorescens*, and *A. faecalis*. Bacteria were cultured under standard aerobic conditions at 30°C in half-strength nutrient broth supplemented with 70 mg/L thiamethoxam for 24 days. Media was sampled at the indicated intervals and residual thiamethoxam levels were measured using HPLC. Control = uninoculated media. Data are expressed as mean (N = 5) ± standard deviation. p<0.0001. Values not connected by the same letters are significantly different.

**TABLE 1.**
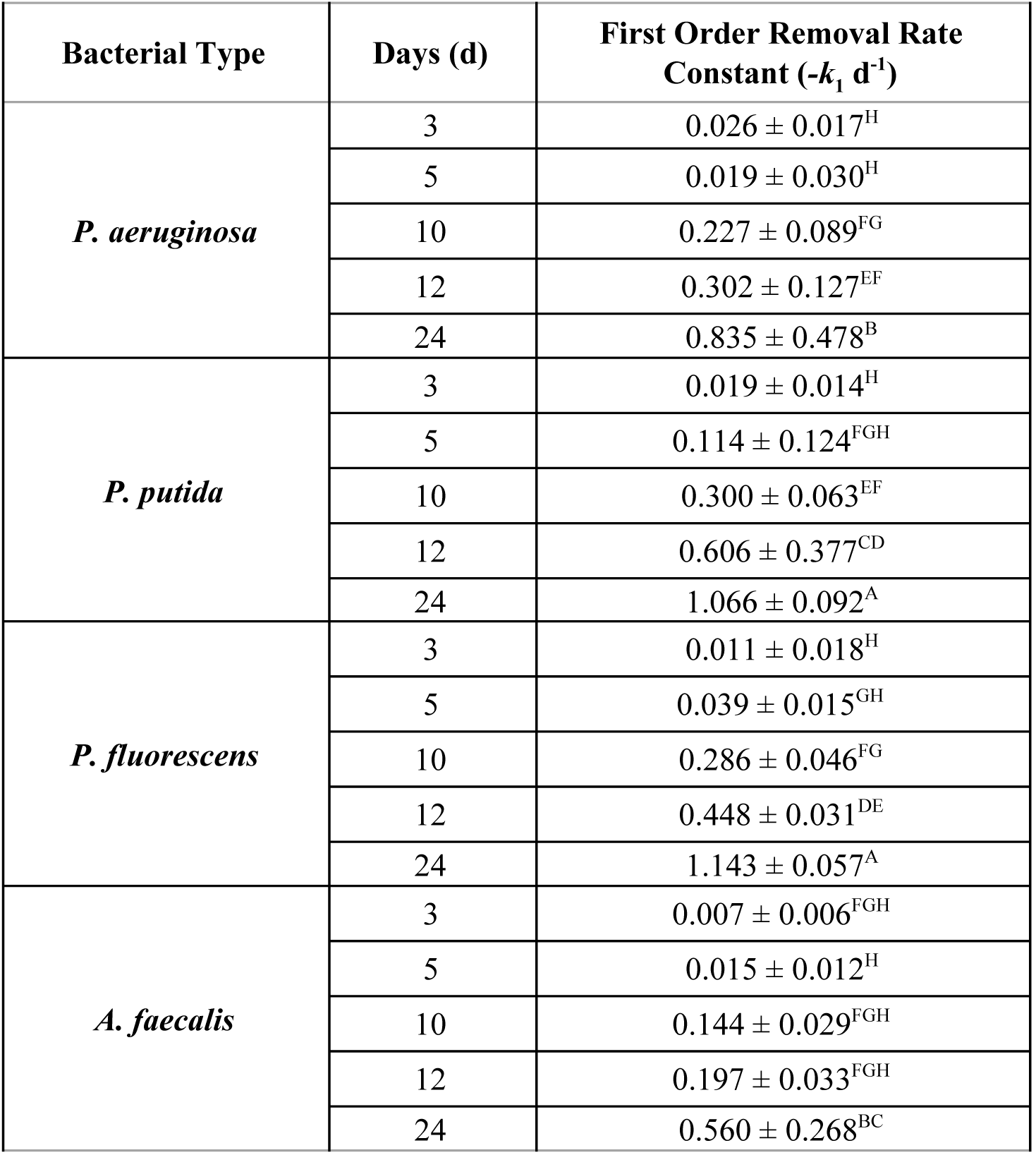
Kinetic parameters for 70 mg/L thiamethoxam removal by *P. fluorescens, P. putida, P. aeruginosa*, and *A. faecalis*. Control = uninoculated media. p<0.001. Values not connected by the same letters are significantly different.

| Bacterial Type | Days (d) | First Order Removal Rate Constant ($-k_1$ d<sup>-1</sup>) |
| --- | --- | --- |
| *P. aeruginosa* | 3 | $0.026 \pm 0.017^H$ |
| | 5 | $0.019 \pm 0.030^H$ |
| | 10 | $0.227 \pm 0.089^{FG}$ |
| | 12 | $0.302 \pm 0.127^{EF}$ |
| | 24 | $0.835 \pm 0.478^B$ |
| *P. putida* | 3 | $0.019 \pm 0.014^H$ |
| | 5 | $0.114 \pm 0.124^{FGH}$ |
| | 10 | $0.300 \pm 0.063^{EF}$ |
| | 12 | $0.606 \pm 0.377^{CD}$ |
| | 24 | $1.066 \pm 0.092^A$ |
| *P. fluorescens* | 3 | $0.011 \pm 0.018^H$ |
| | 5 | $0.039 \pm 0.015^{GH}$ |
| | 10 | $0.286 \pm 0.046^{FG}$ |
| | 12 | $0.448 \pm 0.031^{DE}$ |
| | 24 | $1.143 \pm 0.057^A$ |
| *A. faecalis* | 3 | $0.007 \pm 0.006^{FGH}$ |
| | 5 | $0.015 \pm 0.012^H$ |
| | 10 | $0.144 \pm 0.029^{FGH}$ |
| | 12 | $0.197 \pm 0.033^{FGH}$ |
| | 24 | $0.560 \pm 0.268^{BC}$ |

**TABLE 2.**
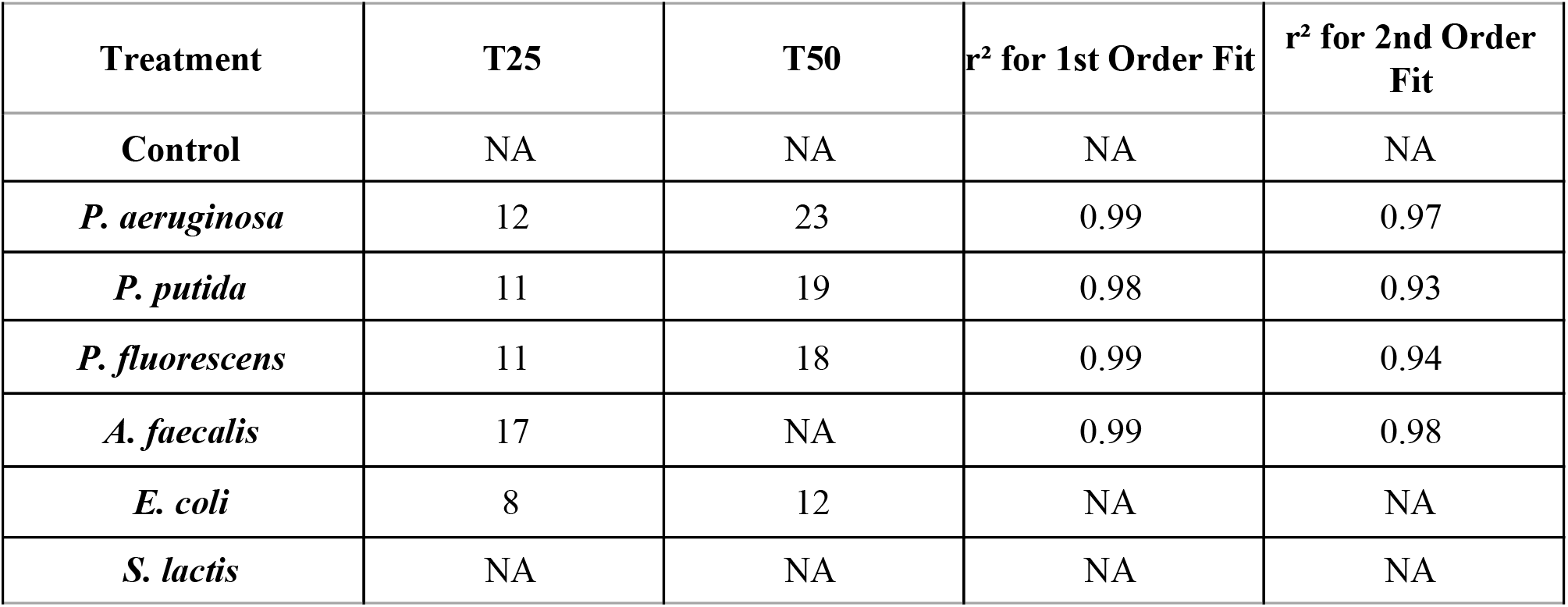
Removal of 70 mg/L thiamethoxam from aqueous solution by *P. aeruginosa, P. putida, P. fluorescens, A. faecalis*, and *E. coli*.

Interestingly, when *E. coli* and *S. lactis* were included in a similar experiment, both showed significant (p<0.0001) reductions in 70 mg/L THM concentration, with *E.coli* exhibiting a 60% reduction and *S. lactis* exhibiting a 12% reduction at the conclusion of the 14-day study period (Fig. 3). When results from Fig. 2 and 3 are taken together, and the organisms’ rates of reduction compared by looking at the amount of time required to reduce 70 mg/L THM concentrations by 50%, *E. coli* notably exhibits the fastest removal capability (T_50_=12 d), followed by *P. fluorescens* (T_50_=18 d), *P. putida* (T_50_=19 d), and *P. aeruginosa* (T _50_=23 d) (Table 2). Neither *A. faecalis* nor *S. lactis* achieved 50% removal during their respective study periods (Table 2).

**FIGURE 3.**
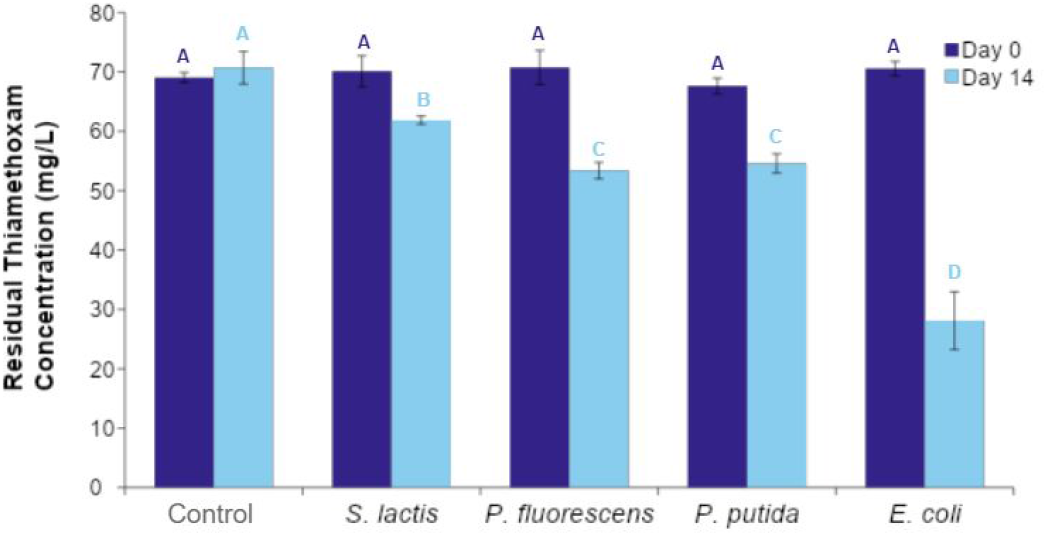
Removal of thiamethoxam from aqueous solution by *S. lactis, P. fluorescens, P. putida*, and *E. coli*. Bacteria were cultured under standard aerobic conditions at 30°C (*S. lactis, P. fluorescens, and P. putidat* or 37°C (*E. colit* in half-strength nutrient broth supplemented with 70 mg/L thiamethoxam for 14 days. Residual thiamethoxam levels were measured after 14 days using HPLC. Control = uninoculated media. Data are expressed as mean (N = 5) ± standard deviation. p<0.0001. Values not connected by the same letters are significantly different.

HPLC analysis of the decrease in THM concentrations over time in the presence of both *P. fluorescens* and *P. putida* revealed the emergence and subsequent significant (p<0.0001) increase in concentration of a presumed metabolite peak, which was first detected on day 7 of the 19-day study (Fig. 4). When this study was repeated with *E. coli*, the same metabolite peak (observed between 3.6 and 3.8 minutes) increased over the course of the 69-day study period, at which time the thiamethoxam peak (observed between 2.8 and 3.2 minutes) became barely detectable (Fig. 5). Comparison of metabolic efficiency as a factor of temperature for *P. fluorescens, P. putida*, and *E. coli* revealed that THM removal was directly correlated with culture temperature, with all three species exhibiting the most removal at 30°C (the highest temperature assessed) (Fig. 6). Notably, at 30°C, THM was almost completely removed by day 50 of the 98-day study for all three species (Fig. 6).

**FIGURE 4.**
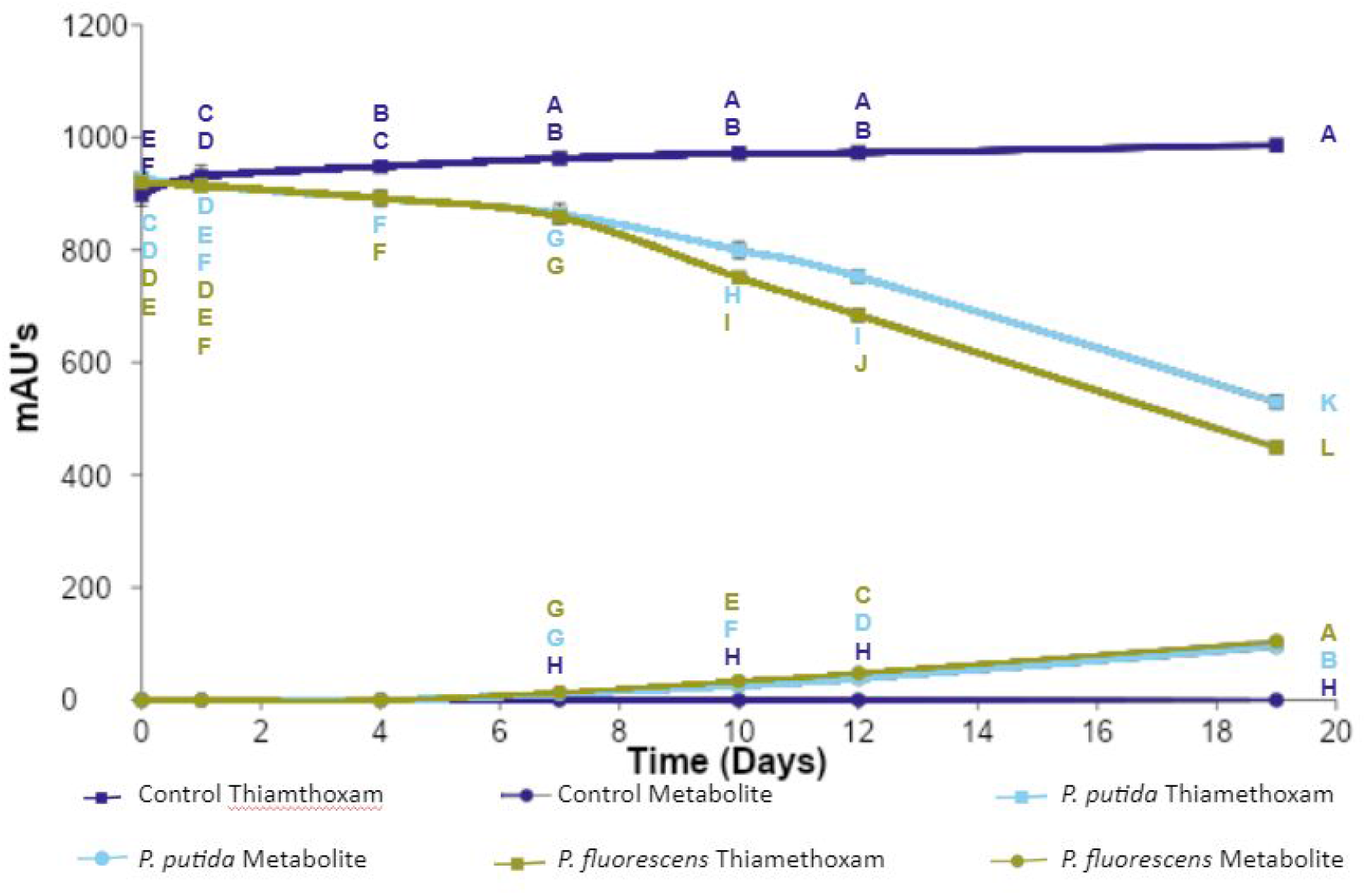
Metabolite generation from thiamethoxam by *P. putida* and *P. fluorescens*. Bacteria were cultured under standard aerobic conditions at 30°C in half-strength nutrient broth supplemented with 70 mg/L thiamethoxam for 19 days. Media was sampled at the indicated intervals and residual thiamethoxam and unidentified metabolite levels were measured using HPLC. Control = uninoculated media. Data are expressed as mean (N=5) ± standard deviation. p<0.0001. Values not connected by the same letters are significantly different.

**FIGURE 5.**
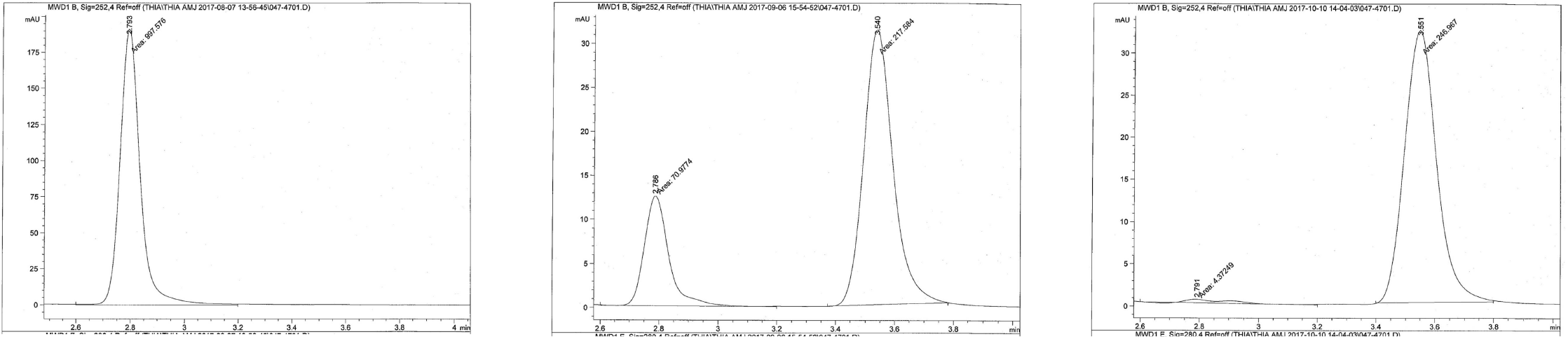
Sample HPLC chromatograms of thiamethoxam and metabolite peaks. *E. coli* was cultured under standard aerobic conditions at 30°C in half-strength nutrient broth supplemented with 70 mg/L thiamethoxam for 98 days. Media was sampled at the indicated intervals and residual thiamethoxam and unidentified metabolite levels were measured using HPLC. Thiamethoxam peaks and metabolites peaks were observed at 2.8-3.2 minutes and 3.6-3.8 minutes, respectively. (A) 14 days, (B) 36 days, and (C) 69 days. Chromatograms are from *E. coli* replicate #3.

**FIGURE 6.**
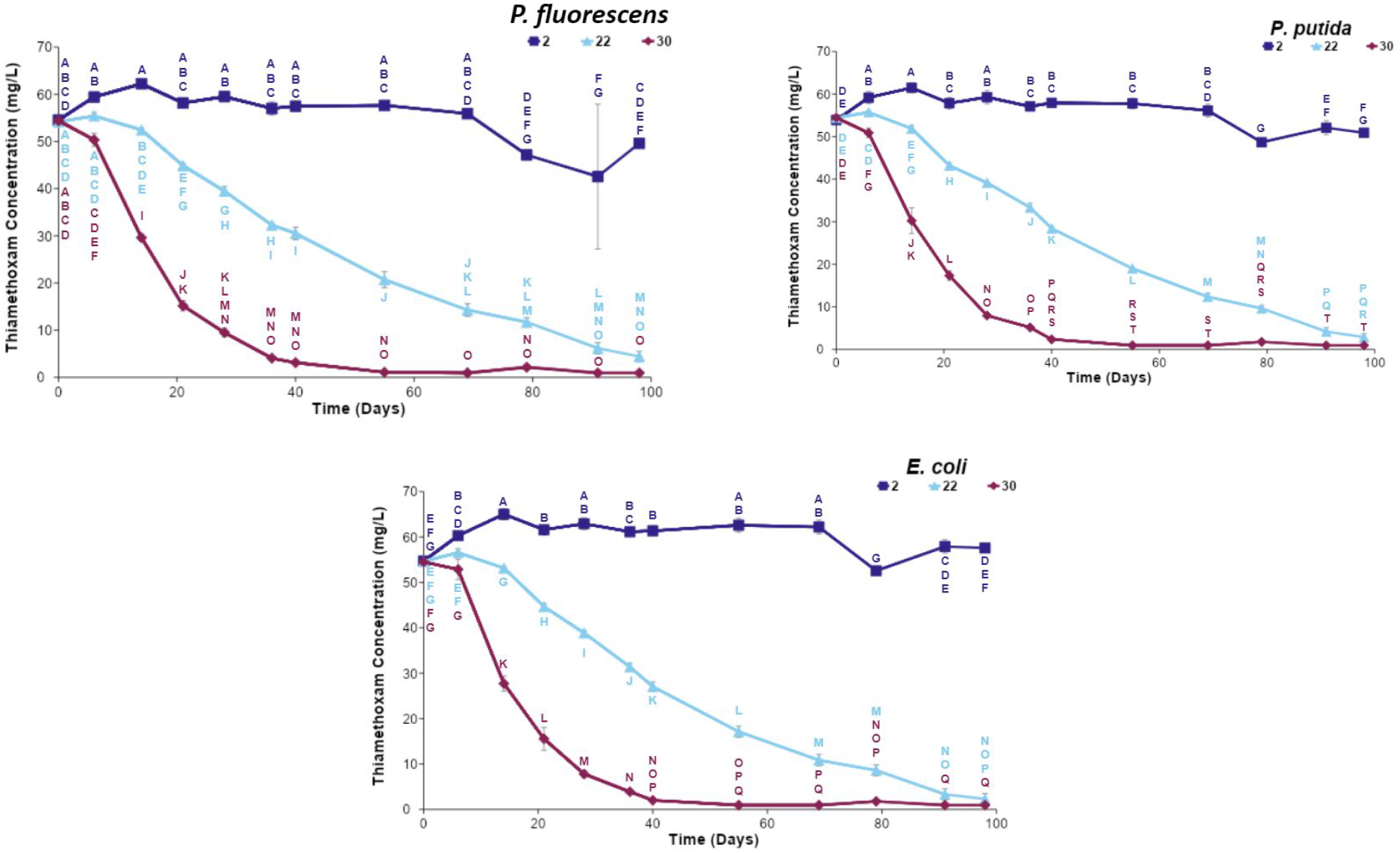
Temperature-dependent removal of thiamethoxam by *P. fluorescens, P. putida*, and *E. coli*. Bacteria were cultured under standard aerobic conditions at 2°C, 22°C and 30°C in half-strength nutrient broth supplemented with 70 mg/L thiamethoxam for 98 days. Media was sampled at the indicated intervals and residual thiamethoxam concentrations were measured using HPLC. Data are expressed as mean (N=3) ± standard deviation. p<0.0001. Values not connected by the same letters are significantly different.

THM removal by *E. coli, P. putida, and P. fluorescens* at 22°C and 30°C also also followed the first order kinetics (see R^2^ in Table 3). All three species achieved the indicated levels of residual THM faster at 30°C than at 22°C, resulting in 95% removal of THM within 38 days for both *E. coli, P. putida*, and 42 days for *P. fluorescens* (Table 3). At 22°C, only *E. coli* showed 95% removal at a much longer time of 82 days whereas *P. putida*, and *P. fluorescens* never reached this level but took 89 and 94 days respectively to attain 90% removal (Table 3). At 30°C after 55 days, the first order rate constant (-*k_1_*/day) was highest in *E. coli*, followed by *P. putida and P. fluorescens*, while at 22°C, the rate constant was not significantly different amongst the species assessed. First order rate constants were determined until THM concentration reached 0 mg/L. The first order rate constants increased over time (Table 4).

**TABLE 3.** Removal of thiamethoxam from aqueous solution by *P. fluorescens, P. putida*, and *E. coli* at 22°C and 30°C. T50, T90, and T95 = time in days required to reduce 70 mg/L thiamethoxam concentration by 50%, 90%, and 95%, respectively.

| Bacterial Species | Temperature (°C) | First Order Removal $R^2$ | T50 | T90 | T95 |
| --- | --- | --- | --- | --- | --- |
| *P. fluorescens* | 22 | 0.97 | 46 | 94 | N/A |
| *P. putida* | 22 | 0.96 | 45 | 89 | N/A |
| *E. coli* | 22 | 0.96 | 42 | 86 | 92 |
| *P. fluorescens* | 30 | 0.99 | 14 | 34 | 42 |
| *P. putida* | 30 | 0.99 | 17 | 36 | 38 |
| *E. coli* | 30 | 0.97 | 15 | 32 | 38 |

**TABLE 4.** First order rate constant of thiamethoxam removal by *P. fluorescens, P. putida*, and *E. coli* at 22°C and 30°C. Bacteria were cultured in triplicate under standard aerobic conditions in half-strength nutrient broth supplemented with 70 mg/L thiamethoxam. Media was sampled weekly and residual thiamethoxam concentrations were measured by HPLC and used to calculate kinetic removal over the study period. p<0.0001. Values not connected by the same letters are significantly different.

| **First Order Rate Constant -k d<sup>-1</sup> at 22°C** |  |  |  |
| --- | --- | --- | --- |
| **Time (days)** | *P. fluorescens* | *P. putida* | *E. coli* |
| **14** | 0.003 ± 0.0005<sup>F</sup> | 0.003 ± 0.0010<sup>F</sup> | 0.001 ± 0.0002<sup>E</sup> |
| **21** | 0.009 ± 0.004<sup>E</sup> | 0.011 ± 0.0006<sup>E</sup> | 0.009 ± 0.0007<sup>DE</sup> |
| **28** | 0.012 ± 0.0009<sup>DE</sup> | 0.011 ± 0.0009<sup>E</sup> | 0.012 ± 0.0001<sup>D</sup> |
| **36** | 0.015 ± 0.0003<sup>CD</sup> | 0.013 ± 0.0009<sup>DE</sup> | 0.015 ± 0.0007<sup>CD</sup> |
| **40** | 0.015 ± 0.0011<sup>CD</sup> | 0.016 ± 0.0004<sup>CD</sup> | 0.017 ± 0.0009<sup>CD</sup> |
| **55** | 0.018 ± 0.0015<sup>BC</sup> | 0.019 ± 0.0001<sup>BC</sup> | 0.021 ± 0.0013<sup>C</sup> |
| **69** | 0.019 ± 0.0013<sup>B</sup> | 0.021 ± 0.0009<sup>B</sup> | 0.023 ± 0.0017<sup>BC</sup> |
| **79** | 0.019 ± 0.0010<sup>B</sup> | 0.022 ± 0.0008<sup>B</sup> | 0.023 ± 0.0018<sup>BC</sup> |
| **91** | 0.024 ± 0.0020<sup>A</sup> | 0.028 ± 0.0026<sup>A</sup> | 0.031 ± 0.0047<sup>AB</sup> |
| **98** | N/A | 0.030 ± 0.0031<sup>A</sup> | 0.033 ± 0.0066<sup>A</sup> |

| **First Order Rate Constant -k d<sup>-1</sup> at 30°C** |  |  |  |
| --- | --- | --- | --- |
| **Time (days)** | *P. fluorescens* | *P. putida* | *E. coli* |
| **6** | 0.013 ± 0.005<sup>D</sup> | 0.010 ± 0.004<sup>E</sup> | 0.004 ± 0.007<sup>E</sup> |
| **14** | 0.044 ± 0.001<sup>C</sup> | 0.042 ± 0.007<sup>D</sup> | 0.048 ± 0.004<sup>D</sup> |
| **21** | 0.061 ± 0.004<sup>B</sup> | 0.054 ± 0.003<sup>C</sup> | 0.060 ± 0.008<sup>CD</sup> |
| **28** | 0.063 ± 0.003<sup>B</sup> | 0.069 ± 0.003<sup>B</sup> | 0.069 ± 0.004<sup>BC</sup> |
| **36** | $0.072 \pm 0.002^A$ | $0.065 \pm 0.001^B$ | $0.073 \pm 0.000^{BC}$ |
| **40** | $0.072 \pm 0.001^A$ | $0.079 \pm 0.001^A$ | $0.082 \pm 0.002^B$ |
| **55** | $0.071 \pm 0.003^A$ | $0.085 \pm 0.001^A$ | $0.104 \pm 0.009^A$ |

## Discussion

Results from the present study indicate that growth of *P. fluorescens, P. putida*, and *P. aeruginosa* in media in which IMI served as the sole carbon or nitrogen source is overall minimal at best. These findings differ somewhat from a number of previous studies which have shown that some *Pseudomonas* strains can metabolize IMI to varying degrees and use it as their sole carbon and nitrogen sources. However, findings across the literature are highly variable, with different groups reporting wide variability in degradative abilities and culture conditions. The differences in our findings are likely due to the different *Pseudomonas* species utilized. Even within the present study, the degradative abilities of the three *Pseudomonas* species studied were quite variable. Additional experiments are underway to follow up with these findings.

Our results show that *P. fluorescens, P. putida*, and *P. aeruginosa* are able to grow in media in which THM serves as the sole carbon or nitrogen source (although growth is substantially less than controls grown in standard media containing a carbon and nitrogen source), indicating that these organisms have the ability to metabolize THM to meet their carbon and nitrogen requirements. These findings align with those of Rana et al. (2015), who found that *P. putida* (strain IMBL 5.2) was able to grow in media in which THM served as the sole carbon and nitrogen source, although, again, growth was less than in standard media. As mentioned previously, this group also found that, despite less growth, THM metabolism was enhanced when it served as the sole carbon and nitrogen source (Rana et al. 2015). However, because these nutrient-limited conditions would be difficult to replicate in the field, further experiments in the present study utilized half-strength nutrient broth, containing both a carbon and nitrogen source.

Our investigations demonstrates that *P. fluorescens, P. putida*, and *P. aeruginosa* are able to degrade THM by 67%, 65%, and 52%, respectively, over a 24-day period, and are consistent with those of Pandey et al. (2009) who identified three *Pseudomonas* strains (1G, 1W, and GP2) capable of degrading THM by approximately 70% over 14 days, and those of Rana et al. (2015) who found that *P. putida* (strain IMBL 5.2) was able to degrade THM by 38% over 15 days. In these experiments, as in the present study, the culture media contained carbon and nitrogen sources in addition to THM.

To our knowledge, the present study is the first one to report the ability of *Escherichia colis* to degrade a member of the neonicotinoid class of insecticides, although *E. coli* has been shown to break down other pollutants, such as the herbicide mesotrione (Olchanheski et al. 2014) and 2,4,6-trinitrotoluene (TNT) (Iman et al. 2017). In the current study, *E. coli* reduced THM levels by 60% over a 14-day period. Notably, *E. coli* exhibited faster degradation of THM than any other species assessed (*E. coli* T_50_=12 d, *P. fluorescens* T_50_=18 d, *P. putida* T_50_=19 d, and *P. aeruginosa* T_50_=23 d). While *E. coli* is known to colonize warm-blooded animals, studies have shown that some strains of *E. coli* are naturalized inhabitants of temperate soils (Ishii et al. 2006). Thus, *E. coli* may represent a novel, highly effective option for the degradation of THM, and may show potential for transformation of other neonicotinoids as well.

Currently, there is a lack of consensus in the literature regarding metabolites generated during THM degradation by *Pseudomonas* species. Results from the present study showing a single prominent metabolite (currently unidentified, observed between 3.6 and 3.8 minutes) generated by *P. fluorescens, P. putida*, and *E. coli* differ from those of Pandey et al. (2009), who detected three metabolites (identified as the desnitro metabolite, retention time 14.05 minutes; the nitrosoguanidine metabolite, retention time 19.65 minutes; and the urea metabolite, retention time 22.72 minutes) generated by *Pseudomonas* (strain 1G), and from those of Rana et al. (2015), who did not detect any metabolites generated by *P. putida* (strain IMBL 5.2). These discrepancies may be due to differences in the organisms, media composition, or HPLC conditions, and underscore the importance of additional research with standardized procedures in this area.

Our results indicate that THM removal capability by *P. fluorescens, P. putida*, and *E. coli* correlates with culture temperature, with all three species exhibiting the highest removal capacity at 30°C (the highest temperature assessed). When cultured at 30°C, THM was almost completely removed by day 50 of the 98-day study for all three species. To our knowledge, the present study is the first to document the kinetic modeling of THM removal and the effects of culture temperature on THM degradation by any microorganism. While one prior study briefly mentioned the first order fit of THM removal only by *P. putida* among twelve tested species (Rana et al. 2014), detailed reports on kinetic modeling with estimated reaction rate constants of THM removal have not been reported thus far.

In conclusion, this work furthers our understanding of *Pseudomonas* species’ ability to degrade THM, documents the emergence of a metabolite that correlates with THM degradation, and identifies *E. coli* as an organism with substantial THM degradation abilities.The findings are promising first step towards the optimization of implementation factors like kinetic parameters and culture conditions for THM transformation by these organisms. Studies are underway to determine the identity of the metabolite and future studies will assess its toxicity relative to THM. If indeed the metabolite exhibits less toxicity than THM, these studies will be repeated in the field. Ultimately, the potential use of these microorganisms as part of a THM bioremediation schema holds promise for a simple, ecologically-friend, sustainable way to remove THM from the environment and help to ameliorate its effects on off-target insects such as honey bees.

## Acknowledgements

The authors wish to thank Grete Bader, Dr. Beverly Brown, Mirzi Devolgado, Sharon Luxmore, Kim Major, Kelsey McNaboe, Alyssa Merrill, Emily Modeen, Jacob Murphy, Janelle Muuse, Gbassey Oteme, Rachel Pacella, Jessica Storrs, and Courtney Taylor for their contributions to this work.

